# Feedback loop regulation between viperin and viral hemorrhagic septicemia virus through competing protein degradation pathways

**DOI:** 10.1101/2024.01.09.574905

**Authors:** Xiaobing Lu, Meisheng Yi, Zhe Hu, Taoran Yang, Wanwan Zhang, E. Neil G. Marsh, Kuntong Jia

**Affiliations:** State Key Laboratory of Biocontrol, Southern Marine Sciences and Engineering Guangdong Laboratory (Zhuhai), School of Marine Sciences, Sun Yat-sen University, Guangzhou, Guangdong, 519082, China; Guangdong Provincial Key Laboratory of Marine Resources and Coastal Engineering, Guangzhou, Guangdong, 519082, China; Departments of Chemistry and Biological Chemistry, University of Michigan, Ann Arbor, Michigan, 48109, USA

**Author notes:** Correspondence: Kuntong Jia,; E. Neil G. Marsh. These authors contributed equally to this work (Xiaobing Lu, Meisheng Yi).

**Keywords:** Autophagy, Immune evasion, Viral hemorrhagic septicemia virus, IRF1, IRF9, Ubiquitination

## Abstract

Viperin is an antiviral protein that exhibits a remarkably broad spectrum of antiviral activity. Viperin-like proteins are found all kingdoms of life, suggesting it is an ancient component of the innate immune system. However, viruses have developed strategies to counteract viperin’s effects. Here, we describe a feedback loop between viperin and viral hemorrhagic septicemia virus (VHSV), a common fish pathogen. We show that *Lateolabrax japonicus* viperin (*Lj*viperin) is induced by both IFN-independent and IFN-dependent pathways, with the C-terminal domain of *Lj*viperin being important for its anti-VHSV activity. *Lj*viperin exerts an antiviral effect by binding both the nucleoprotein (N) and phosphoprotein (P) of VHSV and induces their degradation through the autophagy pathway, which is an evolutionarily conserved antiviral mechanism. However, counteracting viperin’s activity, N protein targets and degrades transcription factors that up-regulate *Lj*viperin expression, interferon regulatory factor (IRF) 1 and IRF9, through ubiquitin-proteasome pathway. Together, our results reveal a previously unknown feedback loop between viperin and virus, providing potential therapeutic targets for VHSV prevention.

**Importance:** Viral hemorrhagic septicaemia (VHS) is a contagious disease caused by the viral hemorrhagic septicaemia virus (VHSV), which poses a threat to over 80 species of marine and freshwater fish. Currently, there are no effective treatments available for this disease. Understanding the mechanisms of VHSV-host interaction is crucial for preventing viral infections. Here, we found that, as an ancient antiviral protein, viperin degrades the N and P proteins of VHSV through the autophagy pathway. Additionally, the N protein also impacts the biological functions of IRF1 and IRF9 through the ubiquitin-proteasome pathway, leading to the suppression of viperin expression. Therefore, the N protein may serve as a potential virulence factor for the development of VHSV vaccines and screening of antiviral drugs. Our research will serve as a valuable reference for the development of strategies to prevent VHSV infections.

## Introduction

The host innate immune response can either be directly activated by viral infection or through the action of interferon (IFN), a key cytokine in innate immunity. IFNs act as master regulators of the immune response and induce the expression of hundreds of so-called IFN-stimulated genes (ISGs) to establish the antiviral state (1). Viperin (virus inhibitory protein, endoplasmic reticulum-associated, IFN-inducible) is among the most highly up-regulated antiviral ISGs (2). It has been found to possess broad-spectrum antiviral activity against DNA and RNA viruses via multiple diverse mechanisms (3). Viperin-like genes have been identified across all 6 kingdoms of life (4, 5), suggesting that it is an ancient defense against viral infection.

Viperin comprises three domains: an N-terminal amphipathic domain, responsible for localizing the protein to the ER and lipid droplets; a central S-adenosyl-L-methionine (SAM) binding domain that contains sequence motifs associated with the radical SAM superfamily, including the canonical CXXXCXXC motif that ligates the catalytic [4Fe-4S] cluster; and the C-terminal domain that binds the substrate, CTP (4, 6). The N-terminal domain is highly variable in sequence and indeed is absent in most viperin-like sequences from microbes (7, 8). In contrast, the SAM-binding and C-terminal domains of viperin are highly conserved across species. Viperin catalyzes the synthesis of the antiviral nucleotide 3ʹ-deoxy-3′,4ʹ-didehydro-CTP (ddhCTP) from CTP. This dehydration reaction occurs by a radical mechanism and is coupled to reductive cleavage of SAM to give 5’-deoxyadenosine and methionine. ddhCTP acts as a chain terminator to inhibit genome replication in some RNA viruses such as Zika virus *in vivo* (9). Recently it has been found that ddhCTP may also induce ribosomal collisions on mRNA resulting in down-regulation of mRNA translation (10).

The transcriptional regulation of viperin appears to be complicated. Various studies have shown that viperin can be induced by both IFN-dependent and IFN-independent pathways during viral infection(11). Viperin is expressed in most cell types at very low levels but can be induced by type I/II/III IFNs (12). By binding with their receptors in either a paracrine or autocrine manner, type I and type III IFNs initiate a signaling cascade that culminates in the formation of signal transducers and activators of transcription (STAT) 1/STAT2/IRF9 complex, termed interferon stimulated gene factor 3 (ISGF3). Then, ISGF3 are translocated to the nucleus to bind the viperin promoter resulting in gene transcription (13). Type II IFN, also named IFN gamma (IFN-γ), is secreted primarily by immune cells and can activate IRF1 to directly induce viperin expression (14). In addition, peroxisomal mitochondrial antiviral signaling protein (MAVS) is responsible for IFN-independent viperin induction by mediating IRF1 activation, leading to it binding the two proximal IRF elements of the viperin promoter and thereby induce viperin expression.

Viperin was initially identified as a protein that was induced by human cytomegalovirus infection or by IFN in fibroblasts, suggesting its antiviral properties (2). Numerous studies point to viperin inhibiting the replication of a broad spectrum of viruses through various mechanisms that remain generally poorly understood (14). Viperin’s original mode of action was likely the inhibition of viral RNA replication through the synthesis of ddhCTP, as noted above. However, viperin has also been shown to binding directly to viral proteins, for example by interacting with the viral protein NS3 viperin suppressed viral genome replication of dengue virus (15, 16). Viperin expression also appears to suppress cholesterol biosynthesis, which inhibits the formation of cholesterol-rich lipid rafts that enveloped viruses such as influenza A need to bud from the plasma membrane of infected cells (17, 18).

ISGs play a vital role in repressing virus infection; however, viruses have developed multiple strategies to evade host elimination and achieve efficient replication by targeting these ISGs. In the case of viperin, it is targeted by the UL41 protein of herpes simplex virus 1 which degrades viperin mRNA to facilitate the sequential expression of viral proteins (19). In a further remarkable example of viral adaption, viperin is “hijacked” by human cytomegalovirus to the mitochondrion where it inhibits fatty acid catabolism (20, 21). The resulting depletion of ATP weakens the actin cytoskeleton facilitating the virus’ exit from the cell. However, apart from these examples, the interplay between viperin and virus remains incompletely understood. In particular, it is unknown if there is a feedback regulatory loop between virus and viperin that maintains immune homeostasis.

In bony fish, viperin was first identified in rainbow trout as a virus-induced gene (vig1) and is highly expressed in VHSV infected leukocytes (22). Although viperins from various fish species have been reported to restrict VHSV infection (23-25), its mechanisms of action remain poorly understood. VHSV is a highly pathogenic fish rhabdovirus belonging to the genus *Novirhabdovirus* of the family *Rhabdoviridae* (26). VHSV infects more than 80 freshwater and marine fishes worldwide and causes severe epidemics of hemorrhagic disease leading to enormous economic losses to the aquaculture industry. The genome of VHSV is a negative-sense, single-stranded RNA molecule (∼11 kb) encoding six proteins: nucleoprotein (N), polymerase-associated phosphoprotein (P), matrix protein (M), glycoprotein (G), nonstructural protein (NV) and large RNA-dependent RNA polymerase (L) (27).

Although several VHSV proteins have been shown to play important roles to affect host immune response during VHSV infection (28, 29), the interplay between VHSV proteins and viperin is still unclear. In this study, the interaction between VHSV and viperin was examined. We demonstrate that sea perch viperin (*Lj*viperin) interacts with and degrades the N and P proteins of VHSV through autophagy to inhibit VHSV replication. This antiviral function of viperin may be evolutionarily conserved. Counteracting this activity, we found that the N protein of VHSV mediates the degradation of IRF1 and IRF9 through the ubiquitin proteasome pathway to block *Lj*viperin expression. This work elucidates previously unrecognized mechanisms utilized by VHSV and viperin to counteract each other, providing valuable insights into potential new therapeutic strategies against VHSV infection.

## Results

### *Lj*viperin is induced through IFN-dependent and independent pathways

Given the diverse nature of the immune system in different vertebrate phyla, we first investigated whether *Lj*viperin could be induced through MAVS and IRF1 activation in fish, as occurs in mammals. The transcription levels of *Ljviperin* and *IRF1* genes were therefore measured in *Lateolabrax japonicus* fry (LJF) cells transfected with MAVS. The results showed that both *Ljviperin* and *IRF1* were up-regulated by MAVS (Fig. 1A). Furthermore, using a luciferase reporter assay, we showed that the IRF1 promoter could be activated by MAVS (Fig. 1B) and then IRF1 activated *Lj*viperin promoter in FHM cells transfected with IRF1 (Fig. 1C). Activation required full-length IRF1, because IRF1 deletion mutants lacking either the N-terminal (IRF1-△N) or C-terminal (IRF1-△C) domains could not activate the *Lj*viperin promoter (Fig. 1D and E). Furthermore, ChIP assay showed that IRF1 could directly bind to the *Lj*viperin promoter (Fig. 1F). These results indicate that viperin in fish is induced through IFN-independent pathways and thus this mechanism is most likely conserved throughout vertebrates.

**FIG 1.**
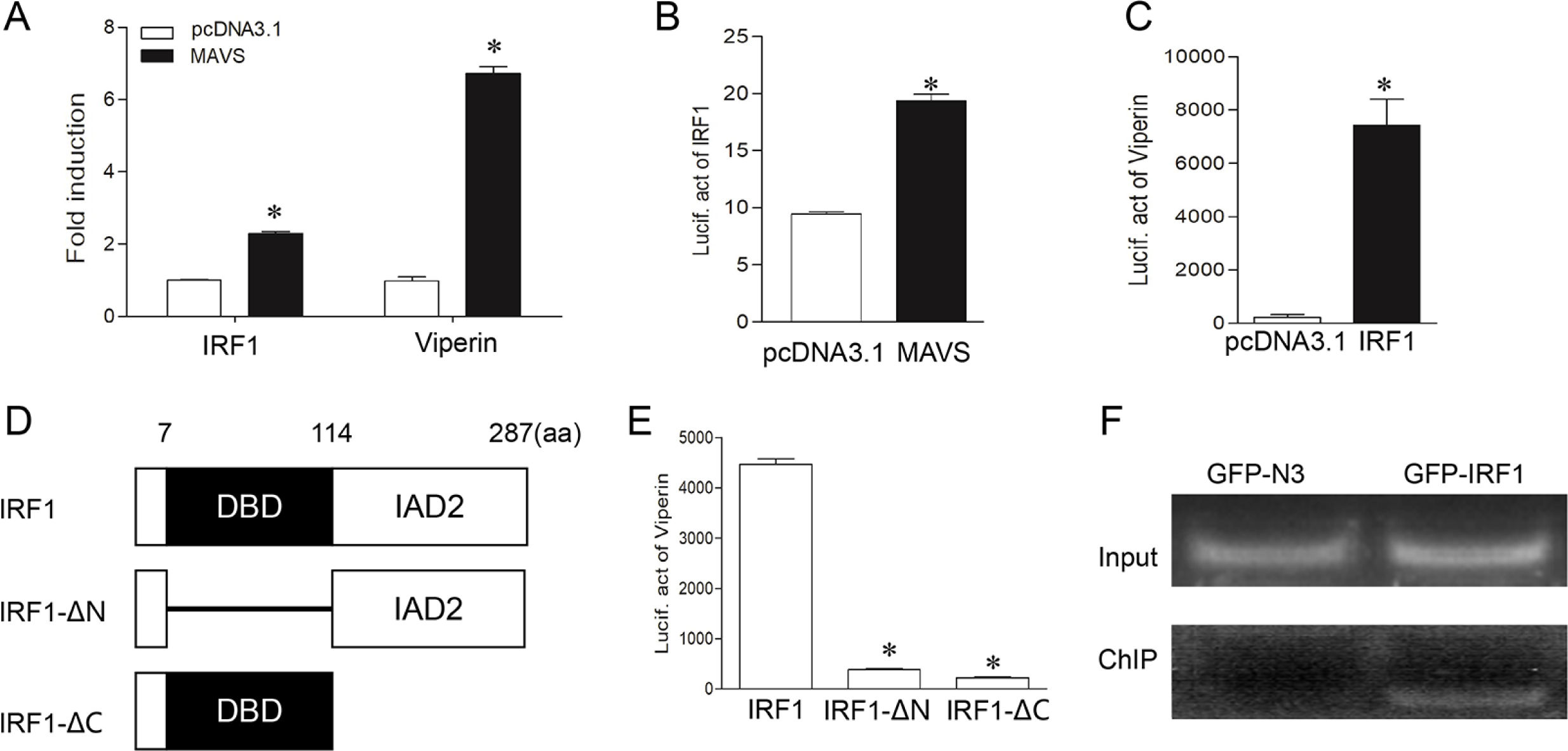
IFN-independent induction of Ljviperin. (A) IRF1 and *Lj*viperin expression are upregulated by MAVS. LJF cells were transfected with MAVS or pcDNA3.1 (empty vector). The mRNA levels of IRF1 and *Lj*viperin were measured by qPCR after 24 h transfection. (B)MAVS activates IRF1 promoter. MAVS was co-transfected with IRF1 promoter in FHM cells. The IRF1 promoter activity was measured by luciferase reporter assay. (C, D and E) Full-length IRF1 is required to induce *Lj*viperin expression. FHM cells were co-transfected with either full-length IRF1 or its domain-deleted variants and the *Lj*viperin promoter constructs. The *Lj*viperin promoter activity was measured by luciferase reporter assay. (F) IRF1 binds to the *Lj*viperin promoter. Either GFP-N or GFP-IRF1 and the *Lj*viperin promoter were co-transfected into HEK 293T cells and the binding of IRF1 to the *Lj*viperin promoter was measured by ChIP assay. All experiments were performed at least three times with similar results. Asterisks indicated statistically significant differences from control (*, *p* < 0.05).

Type I IFNs include IFNc, IFNd and IFNh, whereas type II IFN comprises IFN-γ and IFN-γ rel in sea perch (30). To determine whether the expression of *Lj*viperin was induced by all these IFNs, the transcription levels of *Ljviperin,* together with those of *STAT1*, *STAT2*, *IRF1*, *IRF9*, were measured in LJF cells. As shown in Fig. 2A, IFNc, IFNh and IFN-γ could up-regulate *Lj*viperin expression, but IFNd and IFN-γ rel had no effect on the transcription of *Ljviperin*. Moreover, IFNc and IFNh induced the expression of *STAT1*, *STAT2* and *IRF9*, which then activated *Ljviperin* expression, while IFN-γ mainly induced *Ljviperin* expression through IRF1 activation (Fig. 2B-E). Luciferase reporter assays showed that the *Lj*viperin promoter could only be activated by IRF9, but not by STAT1 or STAT2 (Fig. 2F), suggesting that IRF9 played a key role in the mechanism by which ISGF3 promotes *Lj*viperin expression. Furthermore, we found that the domain-truncated IRF9 mutants (IRF9-△N and IRF9-△C) could not activate *Lj*viperin expression (Fig. 2G and H). To verify whether IRF9 directly activated *Lj*viperin expression, ChIP assays were performed. The results showed that IRF9 could directly bind to the Ljviperin promoter (Fig. 2I). The results suggest that *Lj*viperin can be induced through IFN-dependent pathways, but not all IFN subtypes are capable of up-regulating for *Lj*viperin transcription.

**FIG 2.**
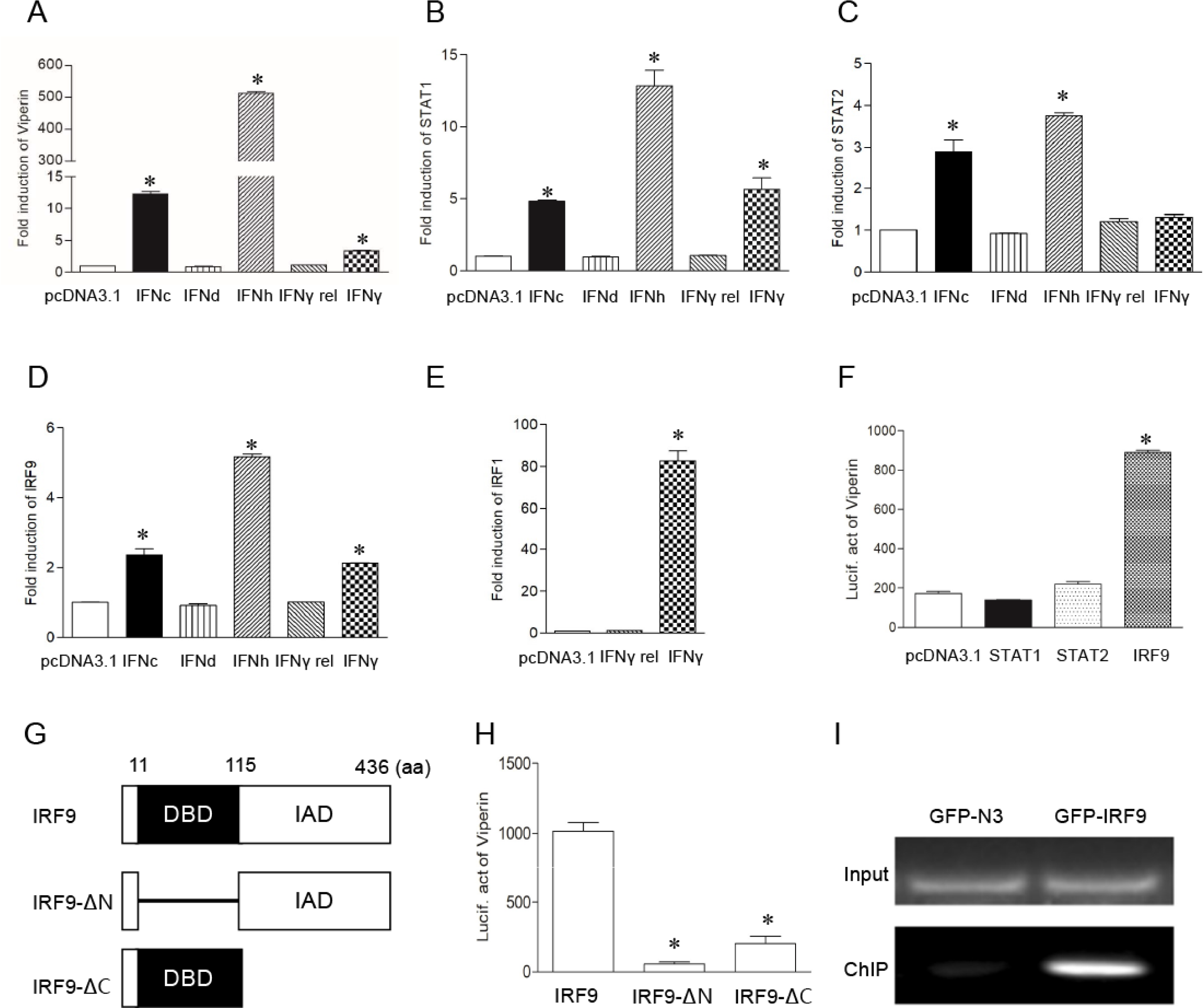
IFN-dependent induction of *Lj*viperin. (A, B, C, D and E) *Lj*viperin is upregulated by various IFNs. LJF cells were transfected with pcDNA3.1 (empty vector control) or the different types of IFNs indicated. The mRNA levels of STAT1, STAT2, IRF9, IRF1 and *Lj*viperin obtained in response to IFN transfection were measured by qPCR. (F, G and H) IRF9 activates the *Lj*viperin promoter. FHM cells were co-transfected with either STAT1, STAT2, IRF9 or domain-deleted variants of IRF9 and *Lj*viperin promoter. *Lj*viperin promoter activity was measured by luciferase reporter assay. (I) IRF9 binds to the *Lj*viperin promoter. GFP-N or GFP-IRF9 and *Lj*viperin promoter were co-transfected into HEK 293T cells and the binding of IRF9 to the *Lj*viperin promoter was measured by ChIP assay. All experiments were performed at least three times with similar results. Asterisks indicated statistically significant differences from control (*, *p* < 0.05).

### *Lj*viperin inhibits VHSV replication

To verify the antiviral activity of *Lj*viperin against VHSV, we first confirmed that it is expressed after VHVS infection in sea perch. The transcripts of *Ljviperin* in spleen were significantly up-regulated and peaked at 24 h after VHSV infection (Fig. 3A). *Ljviperin* expression was also induced after VHSV stimulation *in vitro* (Fig. 3B). Furthermore, overexpression of *Lj*viperin resulted in less CPE compared with the empty vector post-infection with VHSV (Fig. 3C), and significantly inhibited transcription of the viral *N*, *P*, *M*, *G*, *NV* and *L* genes (Fig. 3D). Meanwhile, G protein levels were significantly reduced in cells overexpressing *Lj*viperin (Fig. 3E). We also found that the *Lj*viperin-△DM3 deletion construct lost its antiviral activity, indicating the C-terminal domain was indispensable for *Lj*viperin to suppress VHSV proliferation (Fig. 3F and G). These results show that *Lj*viperin is induced by VHSV infection and inhibits VHSV replication.

**FIG 3.**
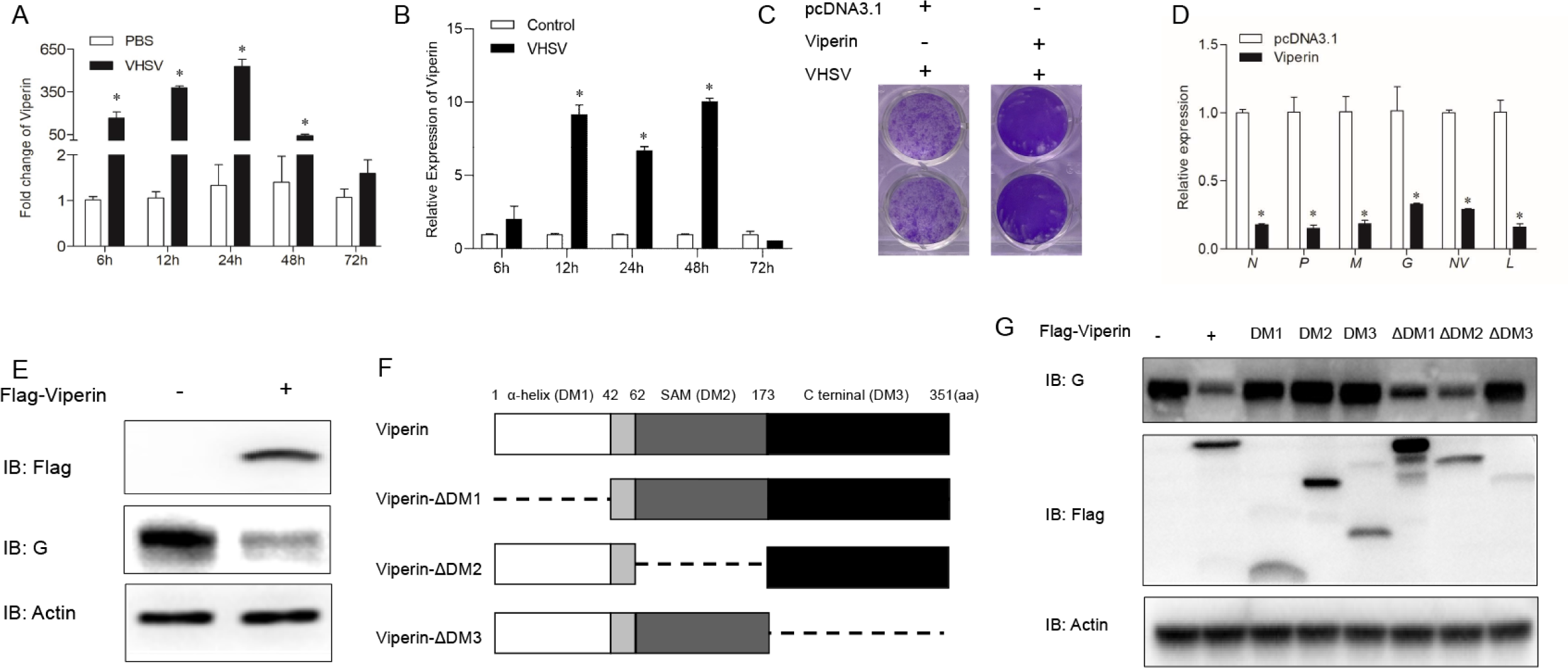
The antiviral activity of *Lj*viperin against VHSV infection. (A) *Lj*viperin expression was measured in sea perch spleen tissue after VHSV infection. The spleen tissues of sea perch were collected at 6, 12, 24, 48 and 72 h after intraperitoneal injection of VHSV, the total RNA was extracted and the mRNA level of viperin was detected by qPCR. (B) *Lj*viperin expression was measured in LJF cells after VHSV infection. LJF cells seeded in six-well plates overnight and then were stimulated with VHSV (MOI=2), the cell samples were collected at 6, 12, 24, 48 and 72 h after VHSV incubation. Gene expression levels of viperin were measured by qRT-PCR. (C) FHM cells seeded in 24-well plates overnight were transfected with 0.5 μg of *Ljviperin* or empty vector. At 24 h posttransfection, cells were infected with VHSV for 36 h. Then, cells were fixed with 4% PFA and stained with 1% crystal violet for visualizing CPE. (D) LJF cells transfected with *Lj*viperin were infected with VHSV; the mRNA levels of the viral *N, P, M, G, NV* and *L* genes were measured by qPCR and compared with non-transfected controls. (E) *Lj*viperin inhibits VHSV replication. Viral G protein levels after infection of *Lj*viperin-transfected FHM cells were measured by IB analysis and compared with empty vector controls. (F and G) Domain-deleted variants of *Lj*viperin were constructed and their antiviral activity against VHSV was analyzed. FHM cells were transfected with full-length *Lj*viperin and domain-deleted variants and then infected with VHSV. Viral G protein expression was detected by IB analysis. Experiments were performed at least three times with similar results. Asterisks indicated statistically significant differences from control (*, *p* < 0.05).

### *Lj*viperin interacts with N and P proteins of VHSV

To investigate which VHSV viral proteins interact with *Lj*viperin, HEK 293T cells were co-transfected with *Lj*viperin and expression vectors for the N, P, M, G NV or L, VHSV viral proteins. The cells were harvested at 24 h after transfection and then the interaction between *Lj*viperin and various viral proteins was examined by Co-IP assays. The results showed that *Lj*viperin bound to both N and P proteins (Fig. 4A-D), but not to M, G, NV, and L proteins (data not shown). Pull-down assays showed that viperin did not directly interact with N or P proteins (Fig. 4E and F), suggesting other proteins may involve in mediating interaction between viperin with N or P protein. We further investigated which viperin domains were necessary to bind the N and P proteins,three *Lj*viperin deletion mutants, respectively containing the α-helix, SAM-binding or C-terminal domains, were co-transfected with N or P proteins in HEK 293T cells and their interaction with the N or P protein was examined by Co-IP. The results showed that the SAM-binding and C-terminal domains of *Lj*viperin interact with the N and P proteins, but the N-terminal domain does not (Fig. 4G and H).

**FIG 4.**
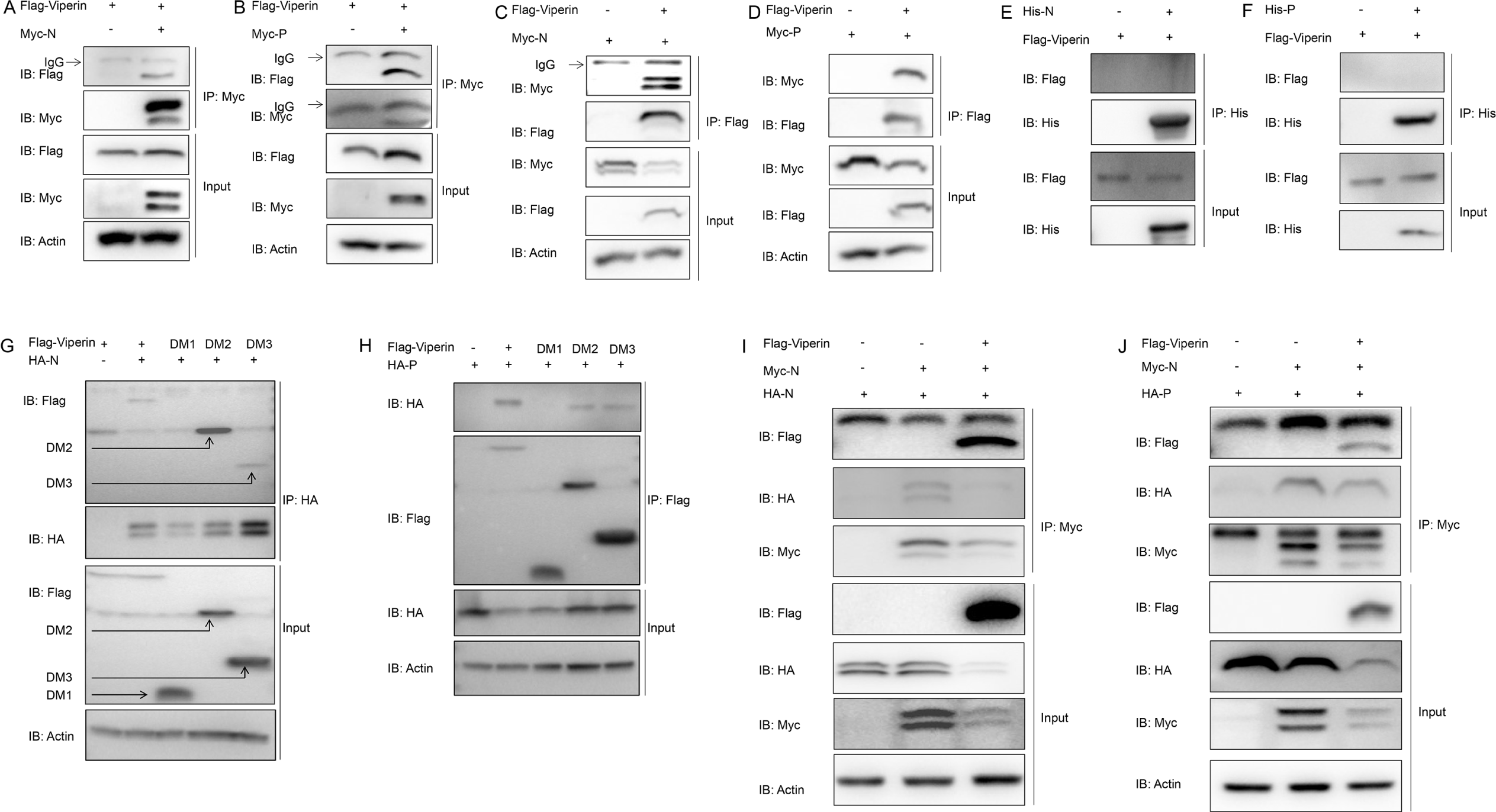
The interaction of *Lj*viperin with viral N and P proteins. (A, B, C, D, G and H) Expression constructs for either the viral N or P proteins and *Lj*viperin or its domain-deleted variants were co-transfected into HEK 293T cells. The interactions of the N or P proteins with *Lj*viperin or its domain-deleted variants were measured by IB analysis. (E and F) The purified His-N proteins and P proteins were incubated with cell lysate prepared from *Lj*viperin-transfected HEK 293T cells, and their interactions detected by IB analysis. (I and J) Homodimerization of N proteins and heterodimerization of N and P proteins were detected in the presence of *Lj*vipeirn. 293T cells were co-transfected with pCMV-Flag or Flag-viperin, Myc-N, and HA-N or HA-P plasmids, respectively. Then the cells were lysed at 24 h post transfection and immunoprecipitated with anti-Myc-conjugated magnetic beads. Western blotting was performed with indicated Abs. All experiments were performed at least three times with similar results.

Homologous dimers of N proteins and heterodimers of N and P proteins of rhabdovirus play important roles in multiple stages of viral replication. To investigate whether *Lj*viperin influences the homodimerization of N proteins and heterodimerization of N and P proteins, Myc-N protein was co-transfected with Flag-Ljviperin and HA-N or HA-P proteins. Co-IP assays found that overexpression of *Lj*viperin significantly reduced the homodimerization of N protein and heterodimerization of N and P proteins (Fig. 4I and J). Taken together, these results suggested that *Lj*viperin binds to the viral N and P proteins through its SAM-binding and C-terminal domains and interferes with the dimerization of these viral proteins.

### *Lj*viperin degrades N and P proteins through the autophagy pathway

As demonstrated in Fig. 4C, 4D, 4I, and 4J, Ljviperin not only interacts with N and P proteins but also appears to degrade them. To further investigate the effect of *Lj*viperin on N and P proteins, Flag-*Lj*viperin and Myc-N or Myc-P expression constructs were co-transfected into FHM cells and HEK 293T cells and then analyzed by immunoblotting. The results showed that *Lj*viperin could degrade N or P protein in a dose-dependent manner in both cells (Fig. 5A and B and Fig. S1A and B). We next explored the pathway by which *Lj*viperin induces degradation of N and P proteins. Flag-*Lj*viperin and Myc-N or Myc-P expression constructs were co-transfected into FHM cells and 293T cells which were then treated with either MG132 or 3-MA, which respectively inhibit the ubiquitin-proteasome and autophagosome pathways. The results showed that *Lj*viperin-mediated degradation of the viral N and P proteins was prevented by 3-MA, but not by MG132 (Fig. 5C and D and Fig. S1C and D). Moreover, 3-MA prevented the degradation of N and P proteins in a dose-dependent manner (Fig. 5E and F and Fig. S1E and F). We also found that the abundance of LC3, which is the marker of autophagy, was enhanced in *Lj*viperin transfected cells (Fig. S2). Interestingly, both human and zebrafish viperins were also found to degrade N and P proteins in a similar manner (Fig. 5G and J and Fig. S1G and J) and inhibit VHSV proliferation (Fig. S3). These results suggest that the degradation viral N and P proteins through the autophagy pathway may be a general feature of viperin’s antiviral activity.

**FIG 5.**
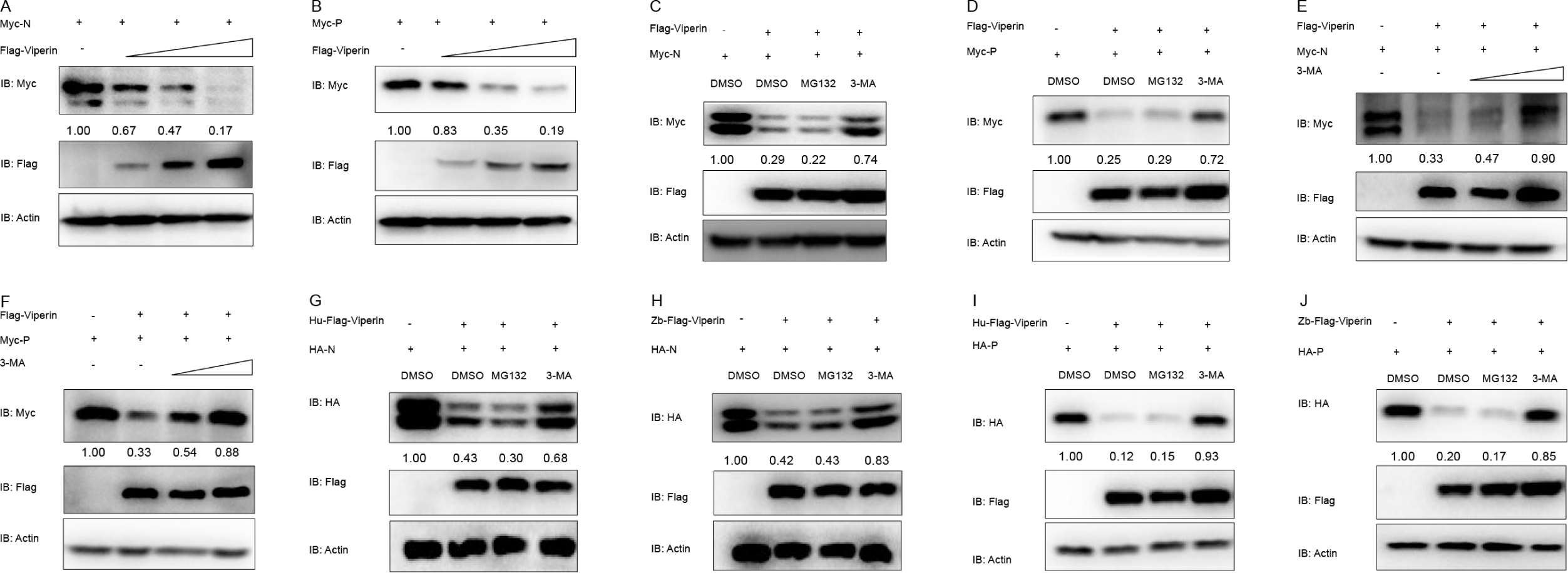
*Lj*viperin degrades viral N and P proteins through the autophagy pathway. (A and B) The abundance of viral N and P proteins decreases in the presence of increasing *Lj*viperin levels. FHM cells were transfected with 2 μg of Myc-N or Myc-P and 2 μg of empty vector, 0.5 μg of Flag-viperin, 1 μg of Flag-viperin, or 2 μg of Flag-viperin. After 24-h transfection, the cell lysates were harvested for IB with the indicated Abs. (C and D) The effect of MG132 and 3-MA on the degradation of viral N and P proteins by *Lj*viperin. FHM cells seeded in six-well plates were transfected with 2 μg of Myc-N or Myc-P and 2 μg of pCMV-Flag or Flag-viperin plasmids, respectively. At 18 h post transfection, the cells were treated with DMSO, MG132 (20 μM) and 3-MA (2 mM) for 6 h. Then, cells were collected for IB with anti-Myc, anti-Flag, and anti-Actin Abs. (E and F) The degradation of viral N and P proteins by *Lj*viperin is rescued by 3-MA in a dose-dependent manner. FHM cells seeded in six-well plates were transfected with 2 μg of Myc-N or Myc-P and 2 μg of PCMV-Flag or Flag-viperin. At 18 h post transfection, the cells were treated with DMSO, 3-MA (2 mM) and 3-MA (4 mM). The cells were collected for IB with anti-Flag, anti-Myc, and anti-Actin Abs. (G, H, I and J) Zebrafish and human viperins degrade viral N and P proteins. Their degradation is prevented by 3-MA. FHM cells seeded in six-well plates were transfected with 2 μg of Myc-N or Myc-P and 2 μg of PCMV-Flag, Hu-Flag-viperin plasmids or Zb-Flag-viperin, respectively. At 18 h post transfection, the cells were treated with DMSO, MG132 (20 μM) and 3-MA (2 mM) for 6 h. Then, cells were collected for IB with indicated Abs. Staining intensity was quantitated by ImageJ software. Numbers beyond the blots represent the relative band intensity of target protein after normalization to β-actin and the band intensity in control cells was set to 1. All experiments were performed at least three times with similar results.

### N protein suppresses *Lj*viperin expression by blocking IRF1 and IRF9 function

Viruses have evolved many strategies to escape the host immune response. To investigate the mechanism by which VHSV evades its immune evasion, we examined the effect of various VHSV viral proteins on *Lj*viperin expression. Luciferase reporter assays showed that the viral N protein significantly antagonized *Lj*viperin promoter activity induced by both IRF1 and IRF9 (Fig. 6A and B). N protein also suppressed expression of endogenous *Lj*viperin induced by poly I:C treatment (Fig. 6C). These results suggested that the N protein of VHSV may affect viperin expression by suppressing the biological function of IRF1 or IRF9.

**FIG 6.**
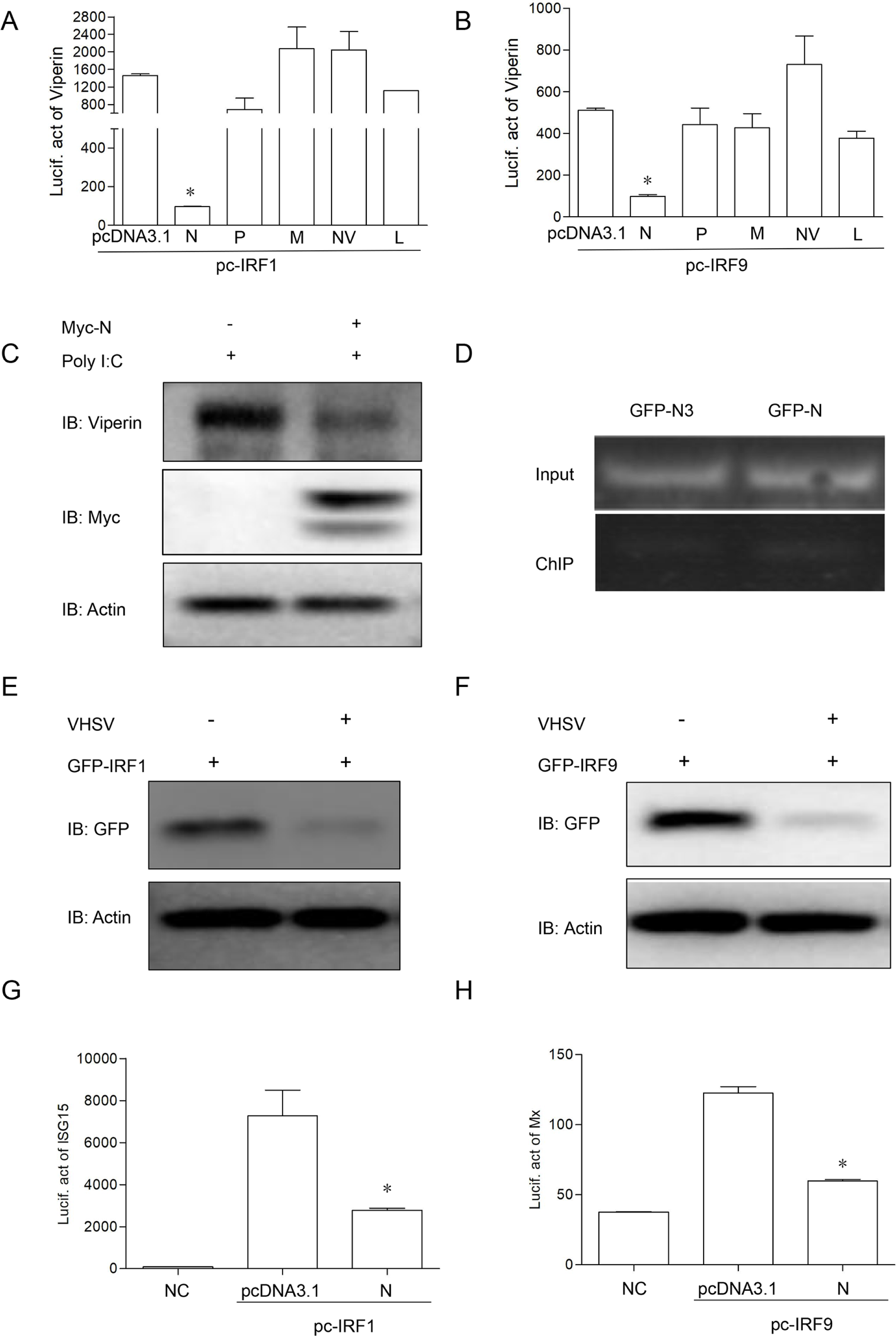
Viral N protein attenuates *Lj*viperin promoter activity by blocking IRF1 and IRF9 function. (A and B) Viral N protein inhibits *Lj*viperin promoter activation induced by IRF1 and IRF9, as determined by luciferase assay. FHM cells were cotransfected with pcDNA3.1, N, P, M, NV or L and IRF1 or IRF9, *Lj*viperin promoter activity was measured by luciferase reporter assay. (C) Viral N protein reduces endogenous *Lj*viperin expresion induced by Poly I:C in LJF cells. LJF cells seeded six-well plates were transfected with 2 μg of PCMV-Myc or Myc-N plasmids, respectively. At 24 h post transfection, the cells were then transfected with 2 μg Poly I:C. The cell samples were collected after 24 h transfection and then analyzed by indicated Abs. (D) The binding of viral N protein with *Lj*viperin promoter. GFP-N3 or GFP-N and *Lj*viperin promoter were co-transfected into HEK 293T cells and the binding of N to the *Lj*viperin promoter was measured by ChIP assay. (E and F) IRF1 and IRF9 are degraded after VHSV infection. FHM cells were transfected with GFP-IRF1 or GFP-IRF9, and incubated with VHSV after 24 transfection. The cell samples were collected after 24 VHSV infection and then analyzed with indicated Abs. (G and H) ISG15 and Mx promoter activation respectively induced by IRF1 and IRF9 are suppressed by N protein. FHM cells were cotransfected with pcDNA3.1 or N and IRF1 or IRF9, *Lj*ISG15 and *Lj*Mx promoter activity were measured by luciferase reporter assay. All experiments were performed at least three times with similar results. Asterisks indicated statistically significant differences from control (*, *p* < 0.05).

To explore whether N protein competitively binds to the *Lj*viperin promoter, GFP-N3 or GFP-N and *Lj*viperin promoter constructs were co-transfected into HEK 293T cells, and cells were harvested for ChIP assays. As shown in Fig. 6D, the N protein does not bind to the *Lj*viperin promoter, suggesting that N protein may block *Lj*viperin expression by affecting either IRF1 or IRF9 expression. Further analysis found that the abundance of both IRF1 and IRF9 was reduced in VHSV infected cells (Fig. 6E and F). The above results suggest that VHSV may utilize N protein to suppress *Lj*viperin transcription by reducing levels of the IRF1 and IRF9 transcription factors. To analyze whether N protein affect other ISGs activated by IRF1 and IRF9, *Lj*ISG15 and *Lj*Mx promoters were co-transfected with IRF1 or IRF9 and pcDNA3.1 or pc-N, the results showed that N protein significantly blocked *Lj*ISG15 and *Lj*Mx promoter activities respectively induced by IRF1and IRF9 (Fig. 6G and H), suggesting that N protein of VHSV may influence the function of many ISGs activated by IRF1 and IRF9.

### N protein promotes ubiquitination of IRF1 and IRF9

To examine the mechanism by which N protein inhibits *Lj*viperin expression, we investigated the interaction between N protein and IRF1 and IRF9. As shown in Fig. 7A, neither IRF1 nor IRF9 were immunoprecipitated by N protein. However, overexpression of N protein reduced the levels of IRF1 and IRF9 in a dose-dependent manner (Fig. 7B and C). Furthermore, N protein-mediated IRF1 and IRF9 degradation was rescued by MG132 in a dose-dependent manner, but not 3-MA (Fig. 7D-G). These data suggest that N protein facilitates degradation IRF1 or IRF9 by targeting them to the proteasome.

**FIG 7.**
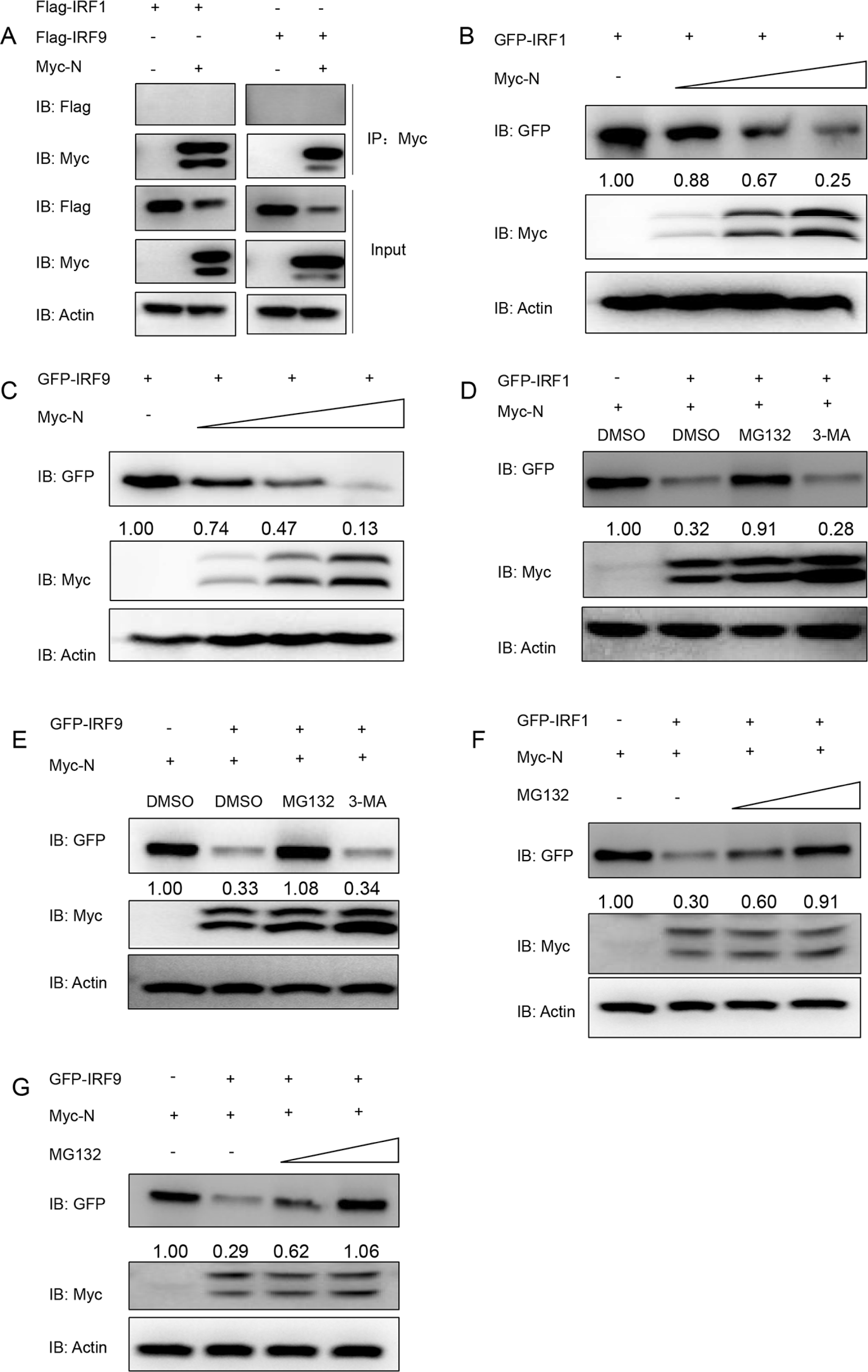
The interaction of viral N protein with IRF1 and IRF9. (A)Viral N protein interacts with IRF1 and IRF9 as determined by IP and IB. 293T cells were transfected with 5 μg of empty vector or Myc-N and Flag-IRF1 or IRF9 plasmids, respectively. At 24 h post transfection, cell lysates were immunoprecipitated with anti-Myc-conjugated magnetic beads and immunoblotted with anti-Myc and anti-Flag Abs; cell lysates were also immunoblotted with indicated Abs. (B and C) Viral N protein degrades IRF1 and IRF9 in dose-dependent manner. 293T cells were transfected with 2 μg of GFP-IRF1 or GFP-IRF9 and 2 μg of empty vector, 1 μg of Myc-N, 2 μg of Myc-N, or 3 μg of Myc-N. After 24-h transfection, the cell lysates were harvested for IB with the indicated Abs. (D and E) The proteasomal inhibitor MG132 inhibits IRF1 and IRF9 degradation induced by viral N protein. 293T cells seeded in six-well plates were transfected with 2 μg of GFP-IRF1 or GFP-IRF9 and 2 μg of PCMV-Myc or Myc-N plasmids, respectively. At 18 h post transfection, the cells were treated with DMSO, MG132 (20 μM) and 3-MA (2 mM) for 6 h. Then, cells were collected for IB with anti-GFP, anti-Myc, and anti-Actin Abs. (F and G) The degradation of IRF1 and IRF9 induced by N protein was prevented by MG132 in a dose dependent manner. 293T cells seeded in six-well plates were transfected with 2 μg of GFP-IRF1 or GFP-IRF9 and 2 μg of PCMV-Myc or Myc-N. At 18 h post transfection, the cells were treated with DMSO, MG132 (10 μM) and MG132 (20 μM). The cells were collected for IB with anti-GFP, anti-Myc, and anti-Actin Abs. Staining intensity was quantitated by ImageJ software. Numbers beyond the blots represent the relative band intensity of target protein after normalization to β-actin and the band intensity in control cells was set to 1. All experiments were performed at least three times with similar results.

To further clarify whether IRF1 or IRF9 was degraded by the ubiquitin-dependent proteasomal pathway, Flag-IRF1 or Flag-IRF9, Myc-N, and HA-Ub were co-transfected into HEK 293T cells. Following immunoprecipitation of IRF1 or IRF9, immunoblot analysis revealed that the N protein promoted the ubiquitination of IRF1 and IRF9 (Fig. 8A). To identify the pattern of ubiquitination of IRF1 or IRF9, HEK 293T cells were co-transfected with Flag-IRF1 or Flag-IRF9, Myc-N and HA-Ub-K27, HA-Ub-K48, or HA-Ub-K63 plasmids. We found that only the Ub-K27 construct supported N protein-promoted ubiquitination of IRF1 and IRF9 (Fig. 8B and C). These results indicate that N protein promotes the K27-linked ubiquitination of IRF1 and IRF9 leading to proteasomal degradation.

**FIG 8.**
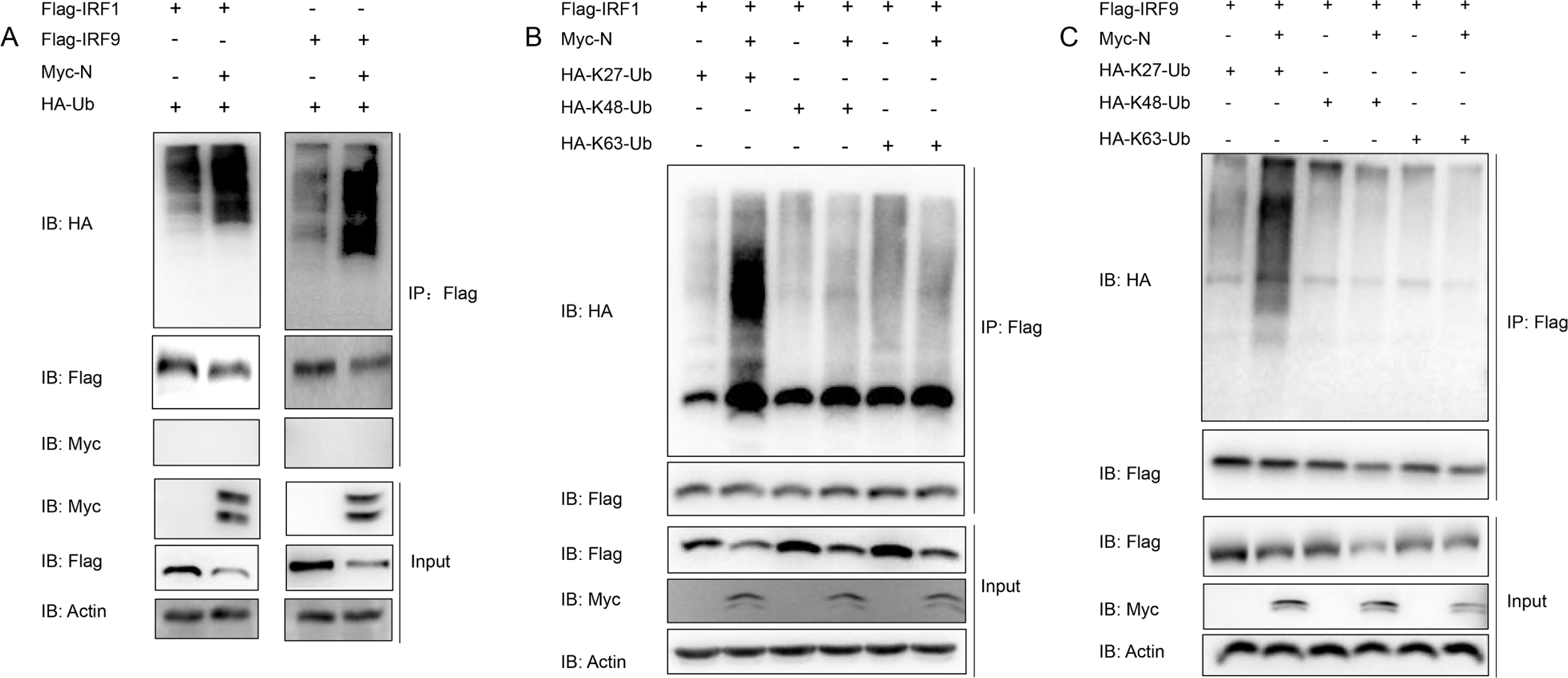
Viral N protein mediates ubiquitination of IRF1 and IRF9. (A) Viral N protein promotes the ubiquitination of IRF1 and IRF9. 293T cells were seeded in 10-cm plates for 24 h and then transfected with 5 μg of Flag-IRF1 or Flag-IRF9, 4 μg of Myc-N or empty vector, and 1 μg of HA-Ub. Cell lysates were immunoprecipitated with anti-Flag magnetic beads and immunoblotted with anti-Flag and anti-HA Abs. The whole-cell lysates were immunoblotted with indicated Abs. (B and C) Viral N protein mediates the K27-linked ubiquitination of IRF1 and IRF9 *in vivo*. Flag-IRF1 or IRF9 was cotransfected with HA-Ub-K27, HA-Ub-K48 or HA-Ub-K63 and Myc-N or pcmv-Myc vectors in 293T cell, the cell samples were collected for IP and then detected by IB analysis.

## Discussion

ISGs are critical for host defense against a diverse group of pathogens. Viperin has attracted particular interest because of its broad-spectrum antiviral activity (14, 31). So far, viperin or viperin-like proteins have been identified in all kingdoms of life and appear highly conserved in both invertebrates and vertebrates (Fig. S4 and S5). This broad conservation points to viperin being an ancient component of the cellular immune response.

In mammals, viperin expression can be induced by IFN-independent and IFN-dependent pathways (32-34). Our studies indicate that the IFN-independent pathway also operates in fish, as we found that MAVS could activate IRF1 expression and thus induce *Lj*viperin expression through IRF1 binding to its promoter. It seems likely therefore that this pathway is conserved in all vertebrate animals, and possibly more broadly. However, whereas in mammals all IFN types induce viperin expression through the formation of the ISGF3 complexes or IRF1 activation (12), we found that in sea perch only two subtypes of type I IFN, IFNc and IFNh, could induce *Lj*viperin expression through ISGF3 activation, and that IRF9 seemed to play the key role in *Lj*viperin induction. The only active type II IFN subtype, IFN-γ induced the expression of *Lj*viperin through IRF1 activation. Multiple IFN subtypes exist in bony fish and our results point to differences between fish and mammals in the IFN-dependent pathway of viperin regulation.

Consistent with studies in other fish and shellfish (23, 25), we found *Lj*viperin was significantly up regulated after VHSV infection *in vivo* and *in vitro*, and overexpression of *Lj*viperin inhibited VHVS proliferation. Interestingly, deletion of the radical SAM domain of *Lj*viperin did not impair the protein’s ability to block VHSV replication. The radical SAM domain is essential for viperin’s enzymatic activity of synthesizing the antiviral nucleotide ddhCTP, which functions as a chain-terminating inhibitor of viral genome replication (9). This observation implies that viperin blocks VHSV replication by mechanisms other than the synthesis of ddhCTP.

The ddhCTP-independent antiviral mechanisms employed by viperin remain poorly understood. However, one common thread is the activation of protein degradation systems. For instance, viperin inhibits replication of hepatitis C virus by promoting the degradation of NS5A through the proteasomal degradation pathway (35). In the present study, we found that *Lj*viperin binds the N and P proteins of VHSV and facilitates their autophagic degradation and thereby blocks VHSV replication. The N and P proteins of mammalian rhabdoviruses play important roles in viral transcription, replication, and assembly (36). Viral RNA is transcribed by the ribonucleoprotein core particles comprising the N, P, and L proteins. Moreover, an N-to-P protein stoichiometry of between 1:1 and 2:1 has been shown to be optimal for efficient vesicular stomatitis virus replication and assembly (37). We found that homodimerization of N proteins and heterodimerization of N and P proteins were significantly suppressed in the presence of *Lj*viperin. Therefore, we speculate that the *Lj*viperin-mediated degradation of N and P may affect the dimerization of these proteins, and thus impede VHSV transcription, replication, and assembly. In addition, rhabdovirus genomic RNA exists as a ribonucleoprotein (RNP) complex, rather than naked RNA, encapsidated with the N protein (38). Degradation of the N protein may expose the RNA genome of VHSV and enable the host intracellular pattern recognition receptors to recognize the viral RNA, which in turn activates the IFN response.

Autophagy is an evolutionarily conserved pathway that targets cellular cytoplasmic constituents for lysosomal degradation, and is essential for maintaining cellular homeostasis in all eukaryotic organisms (39, 40). An increasing number of studies have reported that autophagy can be utilized by ISGs to effectively eliminate of the invading pathogens (41). A recent report found Galectin-9, an IFN-stimulated gene product, cooperated with viperin to restrict hepatitis B virus replication via p62/SQSTM1-mediated selective autophagy (42). Similarly, viperin specifically reduced Ebolavirus VP40 expression by inducing autophagy-mediated lysosomal degradation (43). In this study, we found that *Lj*viperin promoted the degradation of N and P proteins through the autophagy pathway. Further analysis found that N and P proteins interact with the highly conserved SAM and C-terminal domains of *Lj*viperin, however, viperin doesn’t appear to degrade N or P protein directly, suggesting that other proteins are involved in mediating this interaction. So far, these proteins remain to be identified. The observation that both human and zebrafish viperin can mediate the degradation of the N and P proteins through the autophagy pathway suggests that this is an evolutionarily conserved antiviral mechanism.

To survive in their hosts, viruses have also evolved complex mechanisms to evade host antiviral proteins. For example, to escape the antiviral effect of OAS3, an important ISG, enterovirus 71 recruits autologous 3C protease to cleave OAS3 (44). Little is known about how viruses antagonize viperin, but here we uncover a new mechanism by which VHSV suppresses viperin expression. This occurs through the viral N protein targeting IRF1 and IRF9, which are transcription factors that activate viperin expression, and subsequently promoting them for ubiquitination and hence proteasomal degradation. IRF1 and IRF9 are transcriptional activators for multiple genes involved in the innate immune response. It has been reported that ISG15 expression was significantly decreased in IRF1 KO cells (45), and IRF9 can activate the Mx promoter through binding to the ISRE sites (46). Interesting, we found that overexpressed N protein also could inhibit ISG15 and Mx promoter activity respectively induced by IRF1 and IRF9. Therefore, the degradation of these proteins provides viruses with an effective tool to blunt the cellular antiviral response.

The subversion of the protein degradation machinery by viruses to degrade proteins essential for the IFN-mediated immune response is emerging as an important mechanism by which viruses escape the host antiviral response. Other recent examples include the targeting of zebrafish MAVS for K48-linked ubiquitination by the N protein of spring viremia of carp virus, thereby promoting MAVS degradation to block IFN expression (47). Similarly, the ubiquitin-dependent degradation of MITA was promoted by the N protein of hematopoietic necrosis virus to suppress IFN1 production (48). Previously we found that the N protein of VHSV targets STAT1 for ubiquitination and degradation to inhibit MHC-II expression in IFN-γ induced cells and then affect adaptive immunology (49). Therefore, the N protein of VHSV, which affects both host innate and adaptive immune responses, may be a candidate for the development of anti-VHSV drugs in future.

VHSV may lead to acute infection followed by rapid mortality in some infected fish, but it also can establish long term persistent infection in survivors (50). The molecular mechanisms that determine the trajectory of VHSV (either acute or persistent infection) are unknown; however, they likely involve competition between VHSV and host antiviral molecules. Due to its broad-spectrum antiviral activity, modulation of viperin affects both acute and persistent VHSV infections. In the current study, we have uncovered two mutually antagonistic mechanisms by which a) viperin mediates the degradation the viral N and P proteins, and b) the viral N protein suppresses viperin expression.

An interesting consequence of these two opposing positive feedback loops is that they potentially result in a bistable state within the cell that, depending on the VHSV infection process, will either suppress viral replication or, conversely, suppress the antiviral response. Thus, if the cell is primed by IFNs upon VHSV infection so that viperin is already upregulated, then viperin-mediated degradation of N and P proteins will predominate, and virus replication will be suppressed. On the other hand, N protein-mediated degradation of IRF1 and IRF9 will prevent viperin expression, and replication of the virus will be favored with the aggravation of VHSV infection. This will lead to the production of more N protein, reinforcing the degradation of IRF1/9. A third possibility is that neither feedback loop becomes dominant, which would allow a long term, persistent VHSV infection to become established. In this case, we speculate that the complex and mutually antagonistic protein degradation mechanisms employed by VHSV and viperin may lead to a dynamic equilibrium that modulates viperin expression and the ability of VHSV to replicate. One could envision that this balance could be perturbed by various external factors, e.g., temperature change, bacterial infection or other stressors, which would lead to resolution of the infection in favor of either the virus or the cell.

In summary, our studies demonstrate that in bony fish viperin expression is induced by IFN-dependent and IFN-independent pathways. *Lj*viperin targets N and P proteins of VHSV for their autophagic degradation to exert its anti-VHSV function. Given that the sequences of vertebrate viperins are highly conserved and that we showed human viperin is active against VHSV, it seems likely that this is a general mechanism. Through a feedback mechanism, the N protein of VHSV suppresses *Lj*viperin expression by promoting K27-linked ubiquitination and subsequent degradation of the transcription factors IRF1 and IRF9 (Fig. 9). Taken together, these processes represent a novel feedback loop between *Lj*viperin and VHSV, providing fresh insight into the competition between viperin and viruses.

**FIG 9.**
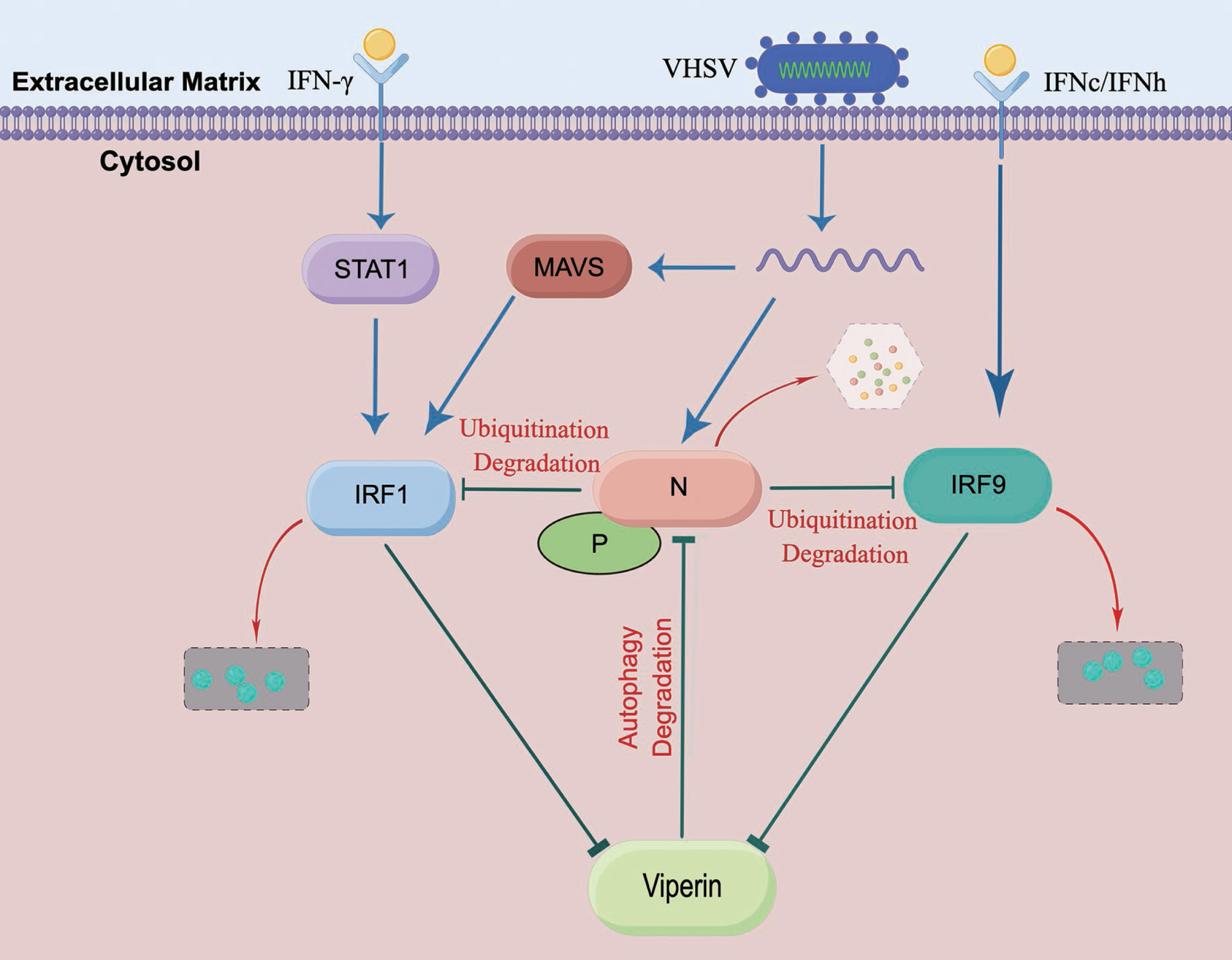
The interaction of viperin and VHSV in bony fish. Viperin is induced through IFN-independent and IFN-dependent pathways and then restricts VHSV infection by degrading the viral N and P proteins through the autophagy pathway. Counteracting this, the N protein of VHSV causes degradation of IRF1 and IRF9 through the ubiquitin-dependent proteasome pathway, there by blocking viperin expression.

## Materials and Methods

### Virus, Cell Culture and Fish

VHSV was isolated from wild largemouth bass *Micropterus salmoides* and kept in our laboratory (26). *Lateolabrax japonicus* fry (LJF) cells and fathead minnow (FHM) cells were respectively cultured at 28°C in DMEM (Invitrogen) and medium 199 (Invitrogen) supplemented with 10% FBS (Life Technologies). HEK 293T cells were maintained at 37°C in DMEM supplemented with 10% FBS. Sea perch (6∼8 cm in body length) obtained from Doumen fish farm (Zhuhai, China) were maintained in recirculating aquaculture system for at least 2 weeks before VHSV infection. All procedures performed with fish were approved by the Ethics Committee of Sun Yat-sen University.

### Genes Cloning, Plasmids Construction and Reagents

IRF1 (accession number OP389271), IRF9 (accession number ON804245), STAT1(accession number OL548908), STAT2 (accession number OP389272), MAVS (accession number KT005247) and viperin (accession number MT437354) sequences of sea perch were obtained from databases derived from published genomic and transcriptome data of sea perch (51). The open reading frames (ORFs) or the mutants of the above genes were cloned into pcDNA3.1 (+), pCMV-Flag or GFP-N3 vectors for overexpression according to experimental requirements. Viperin sequences derived from zebrafish (accession number NM_001025556) and human (accession number NM_080657) were cloned and then inserted into pCMV-Flag vectors. The cDNA fragments of N, P, M, G, NV, and L genes (accession number MK598848) were amplified by RT-PCR and then cloned into pCMV-Myc/pcDNA3.1 vectors, N and P genes were also cloned into pET-23b (+) for prokaryotic expression. For promoter activity analysis, the promoter fragments of *Lj*viperin, *LjISG15* and *LjMx* gene were constructed on basis of pGL3-Basic vector (Promega). All the primers used for vectors construction were shown in Table S1. All constructs were verified by sequencing at TianYi HuiYuan Biological Technology Company (Guangzhou, China). HA-Ub, HA-Ub-K27, HA-Ub-K48, and HA-Ub-K63 plasmids were purchased from YouBia (Guangzhou, China). MG132 and 3-MA were respectively purchased from Sigma-Aldrich and Merck Millipore. Polyinosinic:polycytidylic acid (Poly I:C) were purchased from Sigma-Aldrich and used at a final concentration of 1 mg/ml.

### Cell Transfection and Luciferase Activity Assays

The FHM cells were seeded into 24-well plates overnight and transfected together with various vectors: pcDNA3.1/MAVS/IRF1/STAT1/STAT2/IRF9, IRF1-pro or viperin-pro, and pRL-TK at a ratio of 10:10:1 by lipo8000 Transfection Reagent (Beyotime) according to the manufacturer’s instruction. For the luciferase reporter activity assay, the cell samples were collected after 24 h transfection and washed twice with phosphate buffer saline (PBS), then lysed with Lysis Buffer (Promega) for 15 min on mechanical horizontal rotator, and finally luciferase activity measured using a commercial luciferase assay system (Promega). The assays were carried out independently three times and the average results were used for statistical analysis.

### RNA extraction and quantitative real-time PCR

Total RNAs of spleen tissue or cell samples were extracted by Trizol reagent (Invitrogen) and reverse-transcribed by GoScript reverse transcription system (Promega). The qRT-PCR mixture consisted of 1 µL of cDNA template, 3.5 μl of nuclease-free water, 5 µL of SYBR Green PCR master mix (BioRad), and 0.25 µL of each forward and reverse primers (10 μM). qRT-PCR conditions were as follows: 1 cycle of 5 min at 95°C, 42 cycles of 30 s at 95°C, 30 s at 60°C and 30 s at 72°C, followed by melting curve from 65°C to 95°C to verify the amplification of a single product. Sea perch *β-actin* was used as the internal control. The relative fold changes were calculated by comparison to the corresponding controls using the comparative CT method (2^-ΔΔCt^ method). All primers used for qRT-PCR are shown in Table S1.

### Chromatin Immunoprecipitation Assay

HEK 293T cells were cultured in 10 cm petri dishes and transfected with indicated plasmids. After 24 h, 270 µl of 37% formaldehyde was added to 10 mL of growth media to crosslink intracellular protein-DNA complexes. After incubating at room temperature for 10 min, 0.125 M glycine was added to quench unreacted formaldehyde, and the culture media decanted and discarded. After washing three times with ice-cold PBS, the cell samples were scraped into microfuge tubes and centrifuged at 4°C for 5 min, and then lysed with 1 mL of SDS Lysis Buffer (Beyotime). A quarter of cell lysates were prepared for ultrasonication, and then centrifuged at 4°C for 10 min to remove insoluble material. Sheared crosslinked chromatin (100 µL) was added to the tube containing 900 µL of Dilution Buffer (Sigma-Aldrich). Protein A + G Agarose (60 µL) (Beyotime) was added to pre-clear the chromatin to remove proteins or DNA that may bind nonspecifically to agarose. After centrifugation, 10 µL of the supernatant was removed as input and the remaining supernatant was incubated with anti-GFP-Agarose (Beyotime) overnight at 4°C. Immunoprecipitated GFP-tagged protein-DNA complexes were washed with low salt immune complex wash buffer (0.1% SDS, 1% Tritons X-100, 2 mM EDTA, 20 mM Tris-HCl, 150 mM NaCl), high salt immune complex wash buffer (0.1% SDS, 1% Tritons X-100, 2 mM EDTA, 20 mM Tris-HCl, and 500 mM NaCl), LiCl immune complex wash buffer (0.25 M LiCl, 1% NP40, 1% sodium deoxycholate, 1 mM EDTA, 10 mM Tris-HCl) and TE buffer (10 mM Tris-HCl, 1 mM EDTA) respectively. After washing, 100 µL Elution Buffer (Omega) was added to each tube containing the GFP-tagged protein-DNA complex and then agitated to fully mix with beads. After incubation for 15 min at room temperature, the supernatants were collected into fresh microfuge tubes and the beads were discarded. The samples (immunoprecipitated and input proteins) were treated with 8 µL 5 M NaCl at 65°C for 4-5 h, 1 µL RNase A at 37°C for 30 min, and a solution containing 4 µL 0.5 M EDTA, 8 µL 1 M Tris-HCl and 1 µL proteinase K at 45°C for 2 h. The DNA was purified using a PCR purification kit (Omega). Purified DNA was analyzed by semi-quantitative PCR using specific primers (Table S1).

### Western Blot

Protein samples were separated on SDS-PAGE and transferred to PVDF membrane (BioRad). The membrane was blocked in TBST buffer (25 mM Tris-HCl, 150 mM NaCl, 0.1% Tween 20, pH 7.5) containing 5% skimmed milk for 1 h at room temperature. After incubation with primary antibody (diluted in TBST buffer containing 1% skimmed milk), the membrane was washed three times with TBST Buffer and then incubated with secondary antibody (diluted in TBST buffer containing 1% skimmed milk). After washing three times, the membrane was stained with Immobilon TM Western Chemiluminescent HRP Substrate (Millipore) and examined using an Image Quant LAS 4000 system (GE Healthcare). The antibodies against VHSV G protein, β-actin (Cell Signaling Technology), LC3(Abcam) and viperin (Abcam) were diluted at 1:1000, anti-Flag/Myc/HA/GFP/β-actin (ABclonal Technology) were diluted at 1:2000, and HRP-conjugated anti-rabbit IgG (Beyotime) was diluted at 1:1000.

### Coimmunoprecipitation assay

For coimmunoprecipitation assay experiments, HEK 293T cells were seeded in 10 cm dishes overnight and then transfected with a total of 10 µg of indicated plasmids. At 24 h post-transfection, the medium was removed carefully, and the cell monolayer was washed twice with 10 mL of ice-cold PBS. Then the cells were lysed in 1 mL of radioimmunoprecipitation (RIPA) lysis buffer (Beyotime) containing protease inhibitor cocktail (Sigma-Aldrich). After ultrasonication, cellular debris was removed by centrifugation at 12,000 *g* for 15 min at 4°C. The supernatant was transferred to a fresh tube and incubated with 25 µL of anti-Flag/Myc magnetic beads (MCE) overnight at 4°C with constant agitation. These samples were further analyzed by immunoblotting (IB). Immunoprecipitated proteins were collected by centrifugation at 5,000 *g* for 1 min at 4°C, washed three times with lysis buffer and resuspended in SDS sample buffer. The immunoprecipitated proteins and whole cell lysates were analyzed by IB with the indicated antibodies (Abs).

### Pull-down assay

For prokaryotic expression of His-N or His-P fusion proteins, pET-23b (+)-N plasmids were constructed and then transformed into *E. coli* BL21 (DE3). Bacteria were grown in LB medium at 37°C to an OD_600_ of 0.6∼0.8, and then protein expression induced with 0.4 mM isopropyl-β-D-thiogalactopyranoside at 18°C overnight with agitation at 120 rpm. Cells were pelleted by centrifugation and lysed in lysis buffer (100 mM Na-Phosphate, 600 mM NaCl, 0.02% Tween-20) via ultrasonication. After centrifugation at 12,000 *g* at 4°C for 30 min, the lysate supernatant containing His-tagged proteins was affinity-purified with His-Tag magnetic beads (Invitrogen) and used for pull-down assays. His pull-down assays were performed as described previously (52). Briefly, His-N or His-P-magnetic beads were washed three times with lysis buffer to remove unbound His-N or His-P and were used to bind Flag-tagged protein from the lysates of HEK 293T cells transfected with Flag-viperin. His alone was also prepared and served as a negative control. The beads were washed three times after incubation at 4°C overnight, and then analyzed via immunoblot analysis to detect Flag-viperin proteins.

### Ubiquitination assay

The ubiquitination assay was performed as described previously (53). HEK 293T cells were co-transfected with the indicated plasmids for 24 h. Cells were lysed and then immunoprecipitated with the indicated Abs as described above. The expression of proteins was examined by immunoblotting with indicated Abs.

### Statistics analysis

Statistical analysis was carried out by one-way analysis of variance (ANOVA)with the Dunnett’s post hoc test (SPSS Statistics, Version 20, IBM). All experiments were repeated at least three times.

## Acknowledgments

This work was supported by the National Natural Science Foundation of China (32173001 and 32002417), the Guangdong Natural Science Foundation (2020A1515011169 and 2021A1515110204), Natural Science Foundation of Guangxi Province (2021GXNSFDA075015), China Postdoctoral Science Foundation (2020M683061), Scientific and Technological Planning Project of Guangzhou City (2023B03J1267), Open Project Program of State Key Laboratory of Biocontrol (2023SKLBC-KF03) and the National Institutes of Health (GM 093088 to E.N.G.M.).

## We declare no conflict of interests

Xiaobing Lu and Kuntong Jia designed experiments. Xiaobing Lu, Meisheng Yi, Zhe Hu and Taoran Yang performed experiments, maintained sea perch and cells. Xiaobing Lu and Wanwan Zhang analyzed data. Meisheng Yi and Kuntong Jia supervised the studies. Xiaobing Lu, E. Neil G. Marsh and Kuntong Jia wrote the manuscript.

